# Incorporating sampling bias into permutation tests for niche and distribution models

**DOI:** 10.1101/2022.08.08.503252

**Authors:** Dan L. Warren, Jamie M. Kass, Alexandre Casadei-Ferreira, Evan P. Economo

## Abstract

Randomization tests are often used with species niche and distribution models to estimate model performance, test hypotheses, and measure methodological biases. Many of these tests involve building null models representing the hypothesis that there is no association between the species’ occurrences and the environmental predictors, then comparing the empirical model to null distributions built from these models. These null models are commonly based on points randomly selected with a uniform probability from the study area. However, spatial sampling bias, a near-universal feature of the occurrence data used to build niche and distribution models, results in a non-uniform probability of observing species in different areas even when species occurrences are unrelated to environmental predictors. Failing to account for this bias in randomization tests results in null distributions that do not accurately represent the null hypothesis, potentially leading to incorrect conclusions. In this study, we use simulations to demonstrate that uniform sampling in randomization tests can lead to unacceptable rates of type I error and poor estimates of methodological bias when spatial sampling bias is present in the occurrence data. We present a new method that incorporates a bias estimate into replicate simulations for these randomization tests, and show that this adjustment can reduce type I error rates to an acceptable level.

## Introduction

Species occurrence records from natural history collection data are increasingly employed to answer large-scale biogeographic and ecological questions. The past several decades have seen explosive growth in the development of new methods and computational tools to infer patterns and processes from these data, with a concomitant increase in research output. One major product of this research has been the development of methods for estimating species’ environmental tolerances and potential geographic distributions from occurrence data, termed environmental niche models (ENMs) or species distribution models (SDMs). These models make statistical estimates of species’ environmental associations, often by contrasting conditions at sites where species have been observed with sites without recorded observations, typically represented by a sample of the available area known as “background” data.

While the rapid development of ENMs has produced unprecedented insights into global patterns of biodiversity (e.g., D’Amen et al. 2015; Pineda and Lobo 2009), there are still a great number of methodological issues that limit how trustworthy and interpretable these models are. One of the primary issues with occurrence data is that collection efforts for natural history data are often subject to significant spatial sampling biases. Failing to account for these biases can result in models that parameterize the environmental correlates of sampling bias in addition to, or even at the expense of, estimating species’ environmental tolerances. While methods have been developed that attempt to either reduce the bias in these data sets (e.g., Aiello-Lammens et al. 2015) or correct for bias in the modeling process (e.g., Phillips et al. 2009; Ranc et al. 2017; Vollering et al. 2019; Zurell et al. 2010; Pagel and Schurr 2012; Warren et al. 2021b; Osborne et al. 2022), dealing with spatial sampling bias remains one of the most difficult and confusing aspects of ENM studies.

One of the most common ways that ENM studies attempt to correct for the effects of bias in model construction is to include a bias estimate in the background data (e.g., Mateo et al. 2010; Phillips et al. 2009). With this strategy, background points are sampled from each location with a probability proportional to some estimate of local sampling effort, rather than with uniform probability across the study area. To the extent that this bias estimate reflects the true sampling effort that produced the presence data, this technique is expected to minimize spurious correlations between species occurrences and the environmental predictors due to spatial sampling bias. While in some cases true sampling effort may be known, in many cases it must instead be estimated from data. There are two primary types of data used as bias estimates for background sampling: either occurrence points for closely related species (the “target-group background” approach, Phillips et al. 2009), or points sampled from a grid representing a continuous estimate of spatial sampling bias. These grid estimates are themselves frequently generalizations or derivatives of patterns of sampling in the target group.

The use of any bias estimate necessarily involves making assumptions about the nature of spatial sampling bias, and in many cases these assumptions may have serious implications for interpretations of the resulting models. Broadly speaking, the use of target group data is usually justified by the expectation that the chosen group is sampled using similar approaches to the focal species, and will thus exhibit similar patterns of spatial sampling bias. However, the target group approach also implicitly assumes that spatial sampling bias is the primary phenomenon generating environmental associations that are shared across these groups of species. While this assumption may hold true in some cases, it may be problematic when the target-group species have broad niche overlap with the focal species. The use of target group data is therefore closely tied to the question of niche conservatism: when the environmental niche is largely conserved across the target group and sampling is conducted with no spatial sampling bias, the result is a “niche estimate” for the focal species that only parameterizes the differences between the niche of that species and the shared niche of the target group. This minimizes the effects of conserved aspects of the environmental niche that may in fact be the primary determinants of the focal species’ distribution. This may be particularly problematic in studies where the goal of modeling is to test hypotheses regarding niche conservatism. The ideal target group would therefore be a group of species that shares no environmental responses with the focal species or with each other due to common descent, but which were sampled with the same relative geographic effort as the focal species. The identification of such a group of species is problematic at best in empirical studies, and it is likely that in most cases such a group simply does not exist.

Continuous raster estimates of spatial sampling bias are typically made in one of two ways: either as a kernel density estimate made from target-group data (essentially a smoothed version of the target-group background approach) or as a statistical model of observer behavior that models target group occurrence as a function of predictors that are assumed to affect probability of sampling (e.g., proximity to roads, management status of land). The use of kernel density estimates based on target-group data is subject to the same assumptions as sampling directly from occurrences of the target group, although because this technique uses probabilistic sampling, the resulting background does not follow the spatial pattern of the target-group exactly, which may diminish the impact of some of the issues described above.

Correlative models of spatial sampling bias built using target-group data ideally avoid many of the issues of the target-group background approach by focusing on the aspects of the spatial patterns in the data that are driven by bias per se. However, constructing one of these models necessitates making many decisions that echo those made when modeling the focal species: deciding which predictors are important to describe the spatial pattern of sampling, which families of functional responses are appropriate to fit, etc. Further, if the distribution of the focal species is itself affected by one or more predictors used to model sampling bias, the effects of those predictors in the resulting ENM will tend to be reduced. This may be of particular concern when modeling species whose distributions are strongly impacted by anthropogenic disturbance or human-mediated dispersal.

None of the above is intended as an argument against including estimates of spatial sampling bias in ENM construction; as long as such biases are likely to be present in the data, it is arguably useful to include this information in the modeling process. Instead we want to emphasize that including spatial sampling bias in models involves making assumptions both about observer behavior and the biological process we are trying to model. These assumptions are often not addressed explicitly in the modeling process, and may have serious impacts on the interpretability of resulting models.

Many applications of ENMs involve the use of permutations to test hypotheses or estimate confidence intervals for aspects of the modeling process that are difficult to analyze using parametric statistics. One subset of these tests (hereafter “geographic randomization” tests) involves randomly selecting occurrence points from the study area and fitting a model to those data. This procedure has variously been used to evaluate model performance (Bohl et al. 2019; Raes and ter Steege 2007; Warren et al. 2021a), to measure methodological bias when transferring models (Warren et al. 2021b), or to evaluate whether species’ niches are more or less similar than expected given the set of environments that are accessible to them (Warren et al. 2008; Warren et al. 2021a; Heibl and Calenge 2013).

In the tests that evaluate model performance, this procedure is largely intended to deal with problems stemming from the autocorrelation in various data sources used to build ENMs. Although autocorrelation does not generally bias parameter estimates, it does generate bias in the standard errors of those estimates, which may result in spurious correlations that are misinterpreted as biologically meaningful (Anderson 1954; Hawkins 2012). As a result the expected fit of an uninformative model may have very broad error bars. Although (for example) the AUC of a model built using random data should on average be 0.5 (but see Lobo et al. 2008), values substantially above or below this expectation may be obtained simply by chance. This is particularly common when data sets are small. As originally formulated by (Raes and ter Steege 2007, hereafter “RTS test”), models built using randomly selected points from the study area are evaluated on their discrimination accuracy on the data used to train them. By repeating this procedure many times, users construct an approximate distribution of expected model fits based on the specific combination of sample size, methodology, and study area (with accompanying spatial autocorrelation in environmental predictors) if the occurrence of the species is unaffected by the environment. By comparing the observed fit of the empirical model to this distribution, the investigator can evaluate the support for the hypothesis that the observed fit is a statistical artifact of the study design (including choice of study area, number of occurrence points, choice of predictors, etc.). Bohl et al. (2019) modified this test by measuring how well replicate models based on random points predicted the same withheld validation data used to evaluate the performance of the empirical model, while ENMTools (Warren et al. 2021a) splits both empirical and random data into training and test data sets and compares the ability of the empirical model to predict training and test data to separate null distributions for each (hereafter “EST”, for “ENMTools-style significance test”). Although the differences between these methods may seem small, the result is that they are asking somewhat different questions about model fit. Which (if any) of these is a better measure of model quality is an open question, and may vary based on the intended application of the models. However, resolving this issue is beyond the scope of this study.

Warren et al. (2021b) modified the geographic randomization procedure of Raes and ter Steege (2007) to measure bias in the process of constructing and transferring models. When ENMs are transferred to other times or places, predictions of habitat suitability may be subjected to strong biases specific to a given combination of study area, predictor variables, modeling algorithm, and the time or place to which the models are being transferred. These differences can in some cases result in biases so strong that the occurrence data may have little impact on the prediction being made, which renders their use in decision support questionable. To measure these biases, Warren et al. (2021b) randomly sampled points from the study area, which were then used to construct and transfer models in the same framework as the empirical data, resulting in a set of predictions on the transferred set of layers (e.g., predicted suitability given future climate change) that might be expected by biases related to the study design alone.

Geographic randomization has also been used to test hypotheses of niche evolution. Warren et al. (2008; 2021a) developed a geographic randomization test variously referred to as the “similarity” test or “background” test, in which the overlap between ENM predictions for two species is compared to the overlap between models built with random points from their respective ranges. The goal of this test is to ask whether a pair of species is distributed in environments that are more, or less, similar than expected given the set of available environments in their (potentially allopatric) ranges. Although initially formulated as an “asymmetric” test in which each species’ empirical model was compared to randomized points for the other species, this was extended in the ecospat package (Di Cola et al. 2017) to include the option of a “symmetric” test in which the overlap between the two empirical models was compared to the distribution of overlaps between models built from randomized points for both species.

In all of these tests’ various implementations, geographic randomization is intended to represent the null hypothesis that the occurrence of a species is unrelated to any of the environmental predictors used in the modeling process. The assumption is that if this were the case, the distribution of the species would be expected to be approximately uniform within the study area. However, this process assumes that observer effort is also uniform over the study area. Given the ubiquity of spatial sampling bias in species occurrence data, this is unlikely to be the case for real data. By ignoring the pattern of spatial sampling bias, these null distributions of points therefore most likely underestimate the level of spatial autocorrelation due to spatial sampling bias that would be seen in a typical uninformative data set, which may be problematic for permutation tests. This could result in tests that overestimate model quality (Raes and ter Steege 2007; Bohl et al. 2019; Warren et al. 2021a), underestimate bias in transferring models (Warren et al. 2021b), and mistake shared sampling bias for ecological similarity (Warren et al. 2008).

In a recent study, Osborne et al. (2022) demonstrated a similar method that modified existing tests by building replicate models using random points that were simulated with the same level of spatial autocorrelation seen in the empirical points. They demonstrated that many hypothesis tests that produced statistically significant results when pseudoreplicate points were simulated without spatial autocorrelation instead became non-significant when spatial autocorrelation was included in the pseudoreplicate simulations. This suggests that the existing literature based on these tests may be subject to an unacceptable rate of type I error (falsely rejecting the null hypothesis), and that pseudoreplicate points should more closely mirror the undesirable statistical properties of our occurrence data. However, their implementation treats all spatial autocorrelation in occurrence data as a statistical artifact, which is not necessarily ideal; occurrence data may demonstrate spatial autocorrelation even in the absence of spatial sampling bias and dispersal limitation, simply as a result of individuals responding independently to a spatially autocorrelated environment (Kühn and Dormann 2012; Hawkins 2012). When this is the case, simulating occurrence points with the same level of autocorrelation present in the empirical data may be mistakenly treating much of the ecological signal of habitat suitability as statistical noise, and may overestimate true type I error rates by attempting to factor it out. The method presented in the current study attempts to overcome this issue by focusing directly on one of the primary sources of autocorrelation that is artifactual, namely spatial sampling bias. We suspect that simulating based on an empirical estimate of spatial sampling bias may produce more desirable behavior for a null model, as the empirical correlation between sampling and the environment will be preserved in pseudoreplicate data sets, rather than the overall autocorrelation of the points.

In this study, we conduct a series of simulations to examine the severity of the issue of ignoring the effects of sampling bias in pseudoreplicate data for ENMs while preserving autocorrelation due to natural environmental associations. We focus on scenarios in which the occurrence of the simulated species is unrelated to the environmental predictors, but where there are varying levels and spatial scales of geographic sampling bias. We hypothesize that in this case failing to account for spatial sampling bias will lead to increased type I error for the hypothesis tests regarding niche evolution and model fit (Warren et al. 2008; Warren et al. 2021a; Raes and ter Steege 2007; Bohl et al. 2019), and will lead to poor estimates of methodological bias using the method from bias estimation method Warren et al. (2021b). We then present a new method that includes spatial sampling bias in the generation of random points, rather than overall levels of autocorrelation in the empirical occurrence data as in Osborne et al (2022), and evaluate its effects on type 1 error and estimates of methodological bias.

## Methods

All simulations were restricted to South America (xmin = -92.67, xmax = -33.83, ymin = - 56, ymax = 13.67). Models were built using all 19 bioclimatic predictor variables from the Worldclim 2 dataset at a resolution of 5 arcmin (approx. 10 km at the equator) (Fick and Hijmans 2017). For simulations testing the Warren et al. (2021b) method of measuring methodological bias in transferred models, we projected each model to the same set of variables for the year 2070, using the CMIP5 climate model and representative concentration pathway (RCP) 8.5.

Each species’ range was simulated as a circular buffer around a randomly chosen point from the study area. Since the goal of these simulations was to study the effects of spatial sampling bias on generating spurious results, it was not necessary to simulate any correlation between the suitability of habitat for the species and the environmental predictors. Grid cells with NA values for one or more predictor variables (e.g., cells representing ocean) were set to be outside the species’ range, and as such the number of grid cells falling within a species’ range may vary substantially between simulations with the same radius. To aid in subsequent analyses, we therefore tracked the number of grid cells within each simulated species’ range. We conducted simulations using radii of 250, 500, 1000, 2000, and 4000 km, resulting in simulated species with ranges varying from around 50 to more than 50,000 grid cells.

From these simulated ranges, we sampled occurrence points for each species using three models of spatial sampling bias, hereafter referred to as “uniform”, “density”, and “modeled”. The “uniform” bias model assumes that all grid cells in a species’ range are equally likely to be sampled. For the “density” bias model, we used a kernel density estimate of sampling intensity constructed from a comprehensive target-group dataset. This dataset was composed of occurrence records for all known species of ants from the Global Ant Biodiversity Informatics database (Guénard, Weiser, and Gomez, n.d.), and was originally created for use as a sampling bias layer for a recent global ant biodiversity study (Kass and Guénard et al. 2022). For the “modeled” bias estimate we used these target-group occurrences to fit a logistic GLM that models sampling probability as a function of local human population density (CIESIN 2016), density of roads and distance to nearest road (Meijer et al. 2018), distance to nearest freshwater source (OpenStreetMap Contributors 2017), an index of the level of human modification of local terrestrial systems (Kennedy et al. 2020), and an estimate of the proportion of each grid cell devoted to agriculture (International Food Policy Research Institute 2019). Out of these predictors, human population density and the human modification index had the greatest explanatory power for the target group. Density of roads and distance to freshwater made smaller contributions to the model, while distance to roads and percent of land devoted to crops had little effect. This model was then used to predict relative sampling intensity in each grid cell. We note that for the simulation portion of the current study it is not necessary to decide which of these bias estimates is most realistic; rather, each is separately treated as the “truth” for the purposes of a given set of simulations.

Simulated species’ occurrence data were used to build species distribution models and conduct permutation tests. We conducted simulations to explore the RTS and EST tests (Raes and ter Steege 2007; Warren et al. 2021a), the background/similarity test (Warren et al. 2008; Warren et al. 2021a), and the bias estimation method from Warren et al. (2021b). Each test was conducted using 100 occurrence points per simulated species and 10000 background points sampled from the species’ true range (i.e., the circular buffer from the simulation). As occurrences of the simulated species were unrelated to the environmental predictors, any correlations detected by a given ENM would either be due to chance or as a byproduct of spatial sampling bias. All analyses of simulated data were conducted using Maxent (Phillips et al. 2006) via ENMTools (Warren et al. 2021a) with default settings.

In order to correct permutation tests for spatial sampling bias we modify the existing tests (Raes and ter Steege 2007; Warren et al. 2008; Warren et al. 2021a; Warren et al. 2021b) so that, instead of always sampling occurrences for each permutation with uniform probability across the study area, random points are sampled in proportion to the estimate of relative sampling effort in each grid cell (i.e. the uniform, density, or modeled bias grids). The permutations thus still represent the null hypothesis that the species is uniformly distributed, but allow the possibility that sampling may not be. To evaluate the extent to which spatial sampling bias can affect the outcomes of these tests, we analyze each simulation with and without the inclusion of bias in sampling occurrences for permutations.

## Results

For the RTS test, where model fit is evaluated on training data, we found very high rates of type I error when empirical data were sampled with spatial sampling bias but pseudoreplicate simulations did not include this bias: the null was incorrectly rejected 65.1% of the time when the density bias estimate was used and 68.1% of the time when the modeled bias estimate was used (Figure 3). When data were simulated with uniform sampling and pseudoreplicates were conducted with uniform sampling, type I errors were produced only 1.5% of the time. For the EST test (Warren et al. 2021a) in which model fit is evaluated on randomly withheld test data, type I errors were even higher when pseudoreplicates did not include bias: 86.7% and 84.5% of analyses for density and modeled bias, respectively, contrasted with 5.6% of simulations when the data was sampled without bias. Re-analyzing these simulated data sets with the bias estimate included in pseudoreplicate simulations reduced these errors to much more acceptable levels: 1.5% (density) and 2.6% (modeled) for the RTS method, and 5.8% and 5.3% for the EST method (Figure 3).

**Figure 1.**
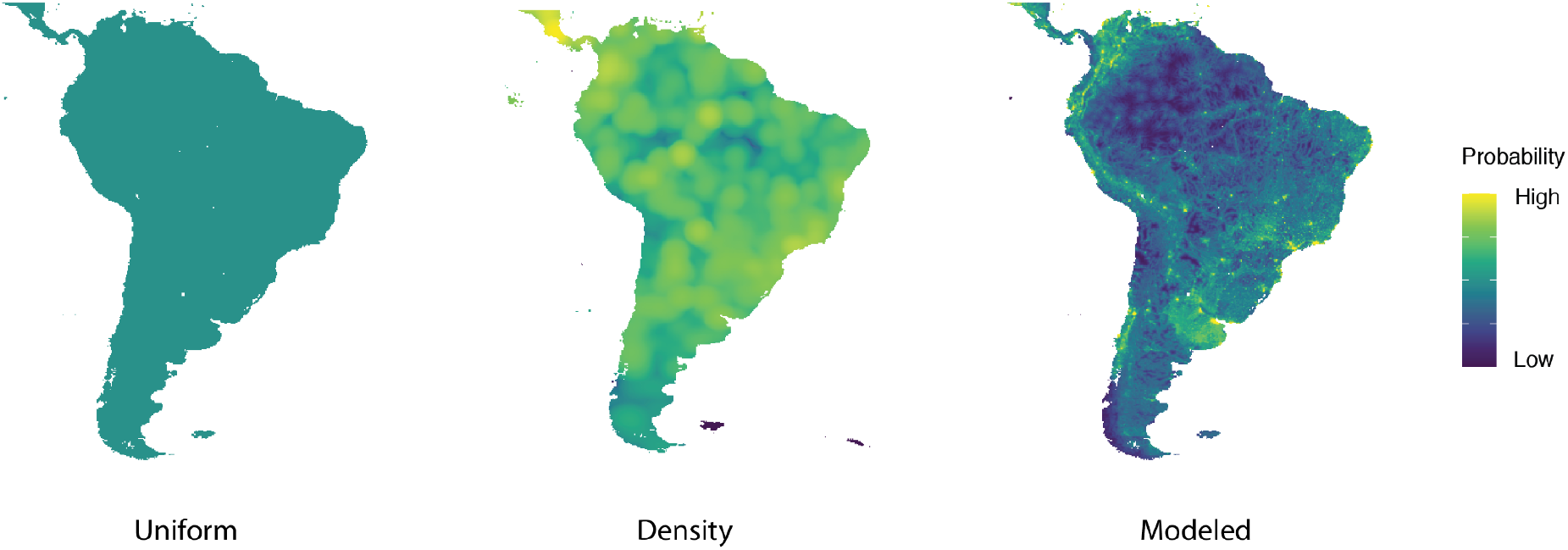
Models of spatial sampling bias. The density bias estimate is based on a kernel density function using target group data, in this case occurrence records for all described ant species from the Global Ants Biodiversity Informatics (GABI) database. The density bias estimate was log transformed prior to plotting to improve visualization of fine scale patterns. The modeled bias estimate is the predicted sampling intensity based on a GLM fitting target group sampling as a function of geographic factors hypothesized to affect sampling.

**Figure 2.**
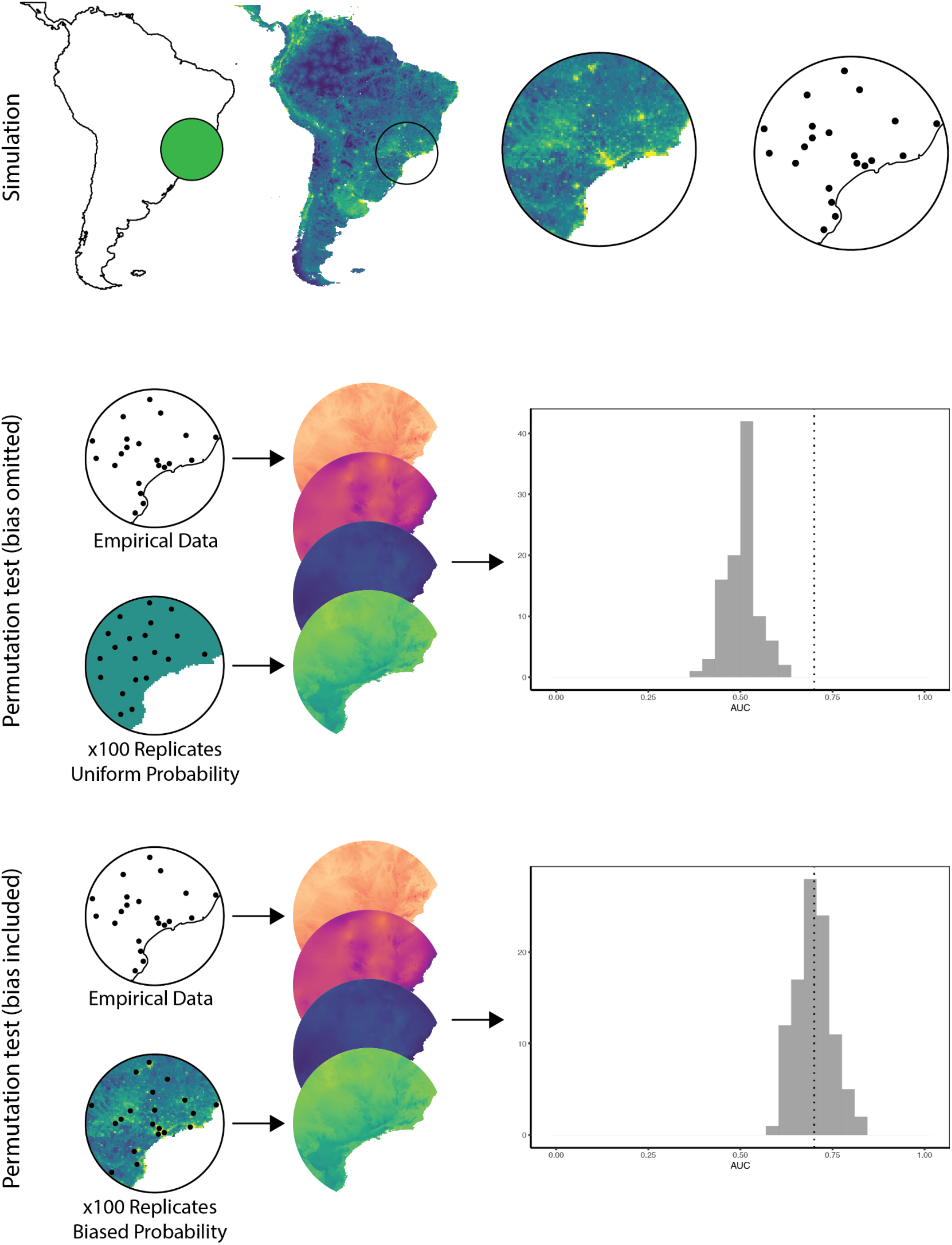
Conceptual outline of simulation and analysis procedures. Simulation (top) begins with placing a circle with a specified radius centered on a random grid cell. This is treated as the species’ range for the purposes of simulation, and raster values for the selected bias model (e.g., GLM modeled bias) are extracted from that range. Points are then sampled in proportion to the sampling intensity in each grid cell determined by that bias model. For analyses, models are built using these simulated points (referred to as “empirical data” for the purposes of analysis) and a set of environmental rasters. When permutations are conducted without including the bias estimate (middle), model performance for the empirical model is compared to a set of 100 models built from points selected at random from the species’ range. As these points are completely uncorrelated with any environmental predictors, performance of these models is generally low, leading us to falsely reject the null (i.e., determine that the model has significant predictive power) because the species’ points are sampled with bias and the permutation points are sampled without that bias. When bias is included in permutation tests (bottom), points for the permutation test are sampled in proportion to the same local sampling intensity that generated the empirical data. This produces a null distribution of expected model performance that is significantly higher than that for uniformly sampled data, leading us to the correct conclusion that performance of the model is no better than expected by chance if the species’ occurrence was unrelated to the environmental predictors.

**Figure 3.**
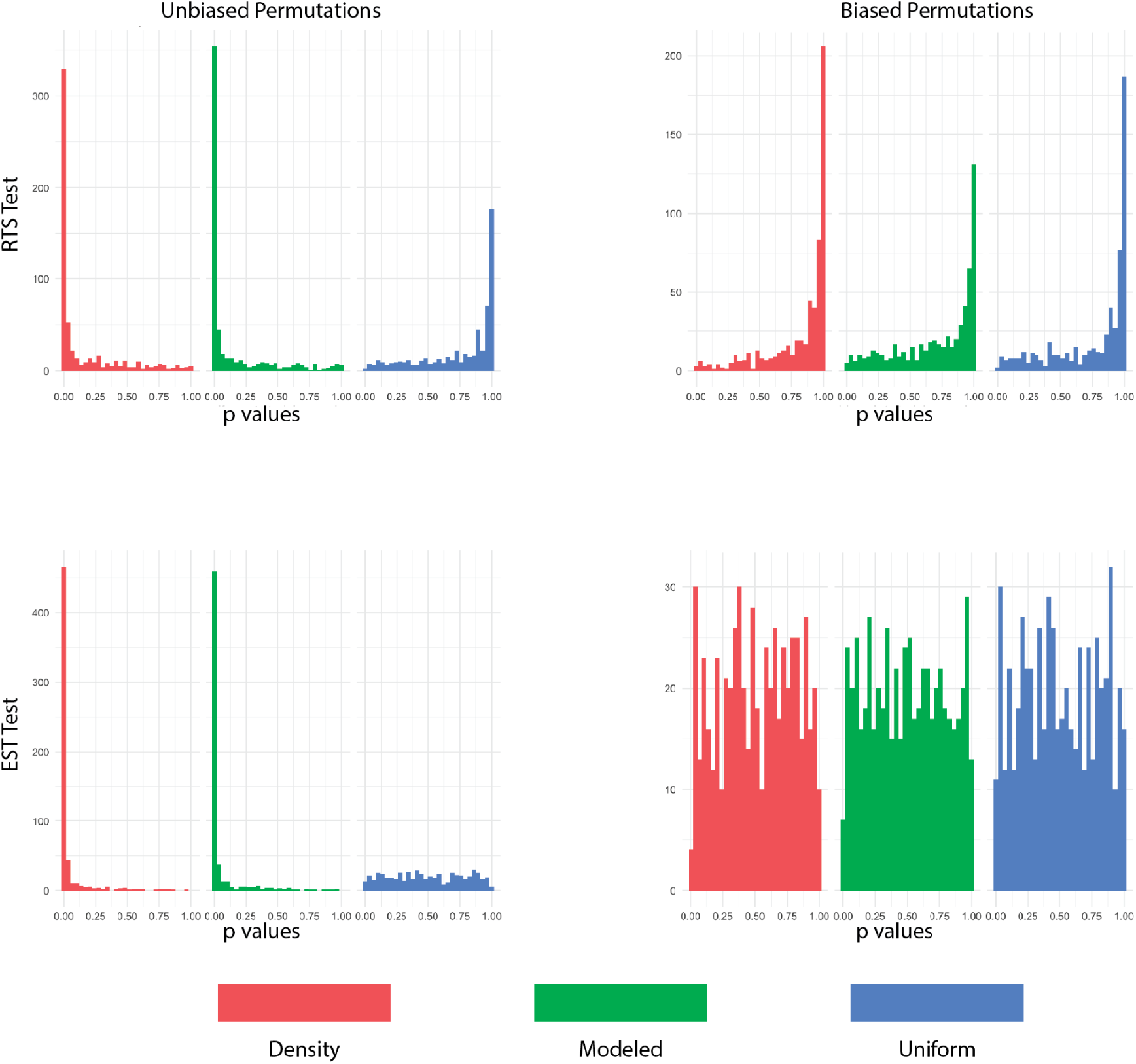
Distribution of p values from Raes and ter Steege (2007, “RTS”) and ENMTools-style (2021, “EST”) tests of model significance, with (right column) and without (left column) inclusion of bias in permutations. Histograms are separated by, and colored according to, the true bias present in the occurrence data. When data is sampled with spatial sampling bias but permutations are conducted without that bias, tests produce a large number of false positives. Inclusion of the bias estimate in the permutations reduces the occurrence of false positives substantially, resulting in a right-skewed distribution of p values for the RTS test and an approximately uniform distribution of p values for the EST test.

For the background tests of Warren et al. (2008), we again found high rates of type I error when data were sampled with bias that was not incorporated into permutations (Figure 4). This was considerably more problematic for the density bias model (type I error rate of 0.42 when overlap was measured using Schoener’s *D*, 0.44 for the *I* metric, and 0.62 for rank correlation) than for the modeled bias estimate (*D*: 0.11, *I*: 0.11, rank correlation: 0.21). The results for the uniform sampling are quite similar to those for the modeled bias estimate, likely because of the spatial scale of autocorrelation for that estimate (*D*: 0.10 *I*: 0.08 rank correlation: 0.09).

**Figure 4.**
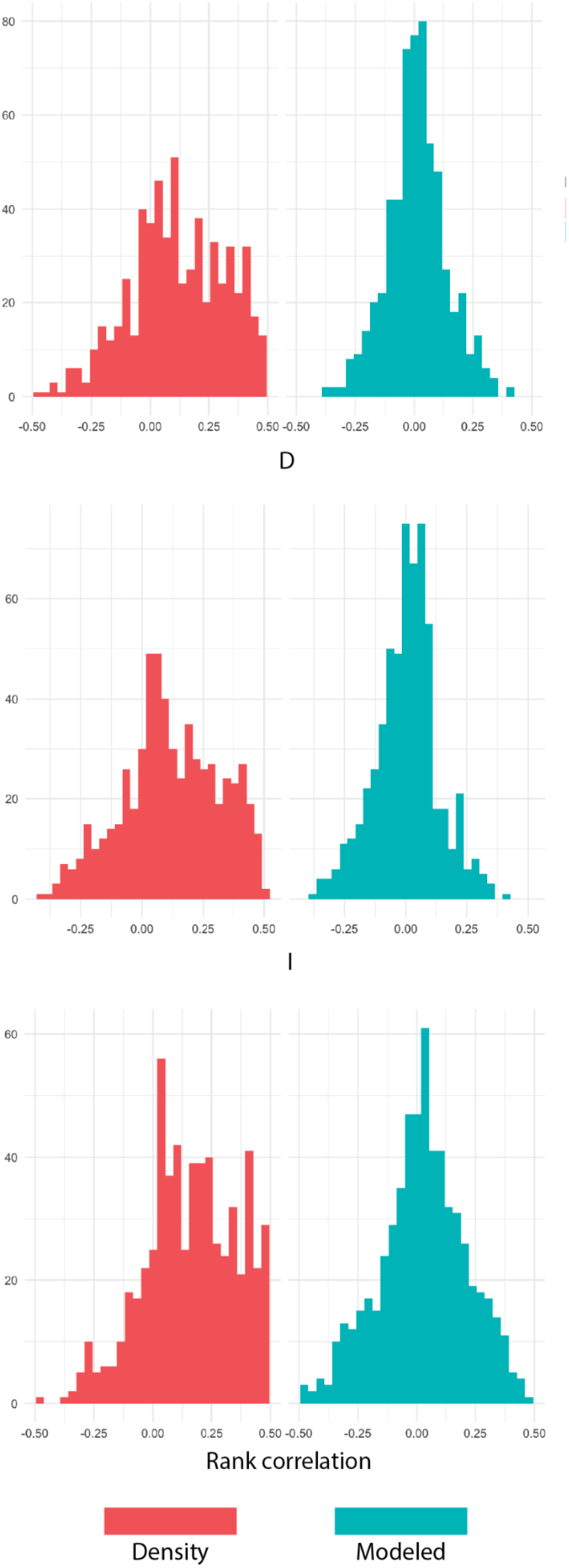
Change in p values from including spatial sampling bias in permutations for the background/similarity test. Positive values indicate increased p value when spatial sampling bias is included. Histograms are colored by the bias model used to generate data and permutations. Results are presented for tests using three different measures of overlap, “D” (top panel), “I” (middle panel), and rank correlation (bottom panel).

For the bias estimation method of Warren et al. (2021b), the goal of permutations is to estimate bias rather than test a hypothesis. As such, type I error rates are not necessarily meaningful to measure. However our results demonstrate that failing to account for spatial sampling bias in the permutations may affect estimated model biases by a substantial margin; average estimated change in habitat suitability across all grid cells may differ by up to 0.6 in either direction, and the number of cells predicted to be declining in suitability in some cases differed by tens of thousands between estimates made from uniformly sampled pseudoreplicates and those sampled including spatial sampling bias (Figure 5). Interestingly, the differences between biased and unbiased pseudoreplicates were generally symmetrically distributed, so that failure to include bias in pseudoreplicates resulted in overestimating future habitat suitability and habitable area as often as underestimating those quantities.

**Figure 5.**
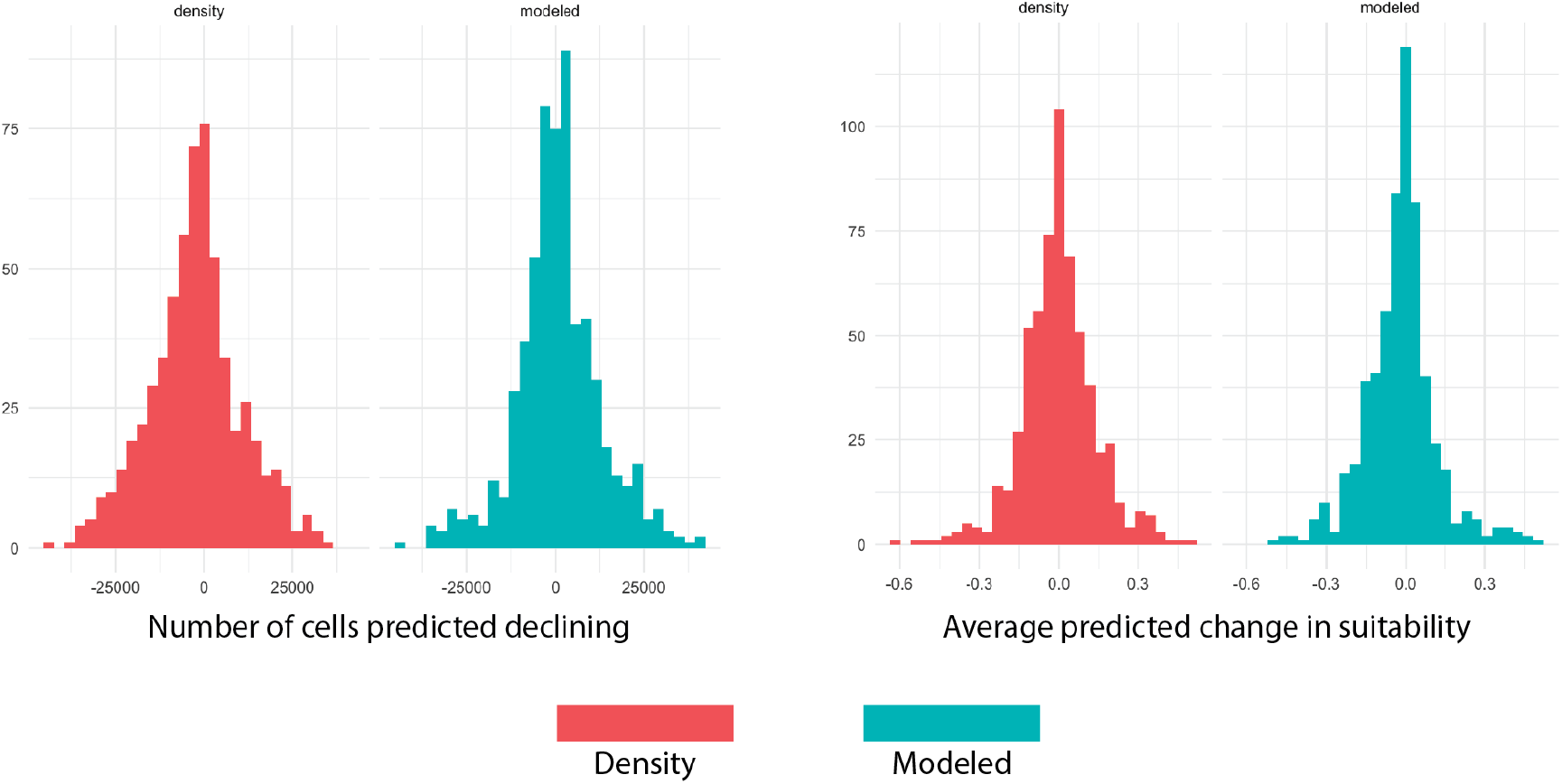
Change in estimates of methodological bias between randomization tests including spatial sampling bias and those without. Left panel depicts change in estimated methodological bias in the number of grid cells predicted to be declining in habitat suitability between the present and future climate scenarios, while the right panel depicts the change in estimated methodological bias in predictions of the magnitude of future change in habitat suitability.

## Discussion

Both the present study and that of Osborne et al. (2022) demonstrate that failure to include non-target processes that affect occurrence data (e.g., spatial autocorrelation or sampling bias) in randomization tests results in unacceptable levels of type I error. In order to correct for this issue, Osborne et al. (2022) modified existing geographic randomization tests by simulating replicate occurrence points with a level of spatial autocorrelation derived from the empirical data set. They showed that many tests on empirical data that showed statistically significant results when replicate data was sampled uniformly were no longer significant when spatial autocorrelation was included in occurrence data for replicates. This is a significant step forward for these tests, but may not be suitable for every study, as it assumes that all spatial autocorrelation is a statistical artifact that must be eliminated.

For that reason we chose in this study to focus on the effects of spatial sampling bias directly. The results of our simulation largely support the conclusions of Osborne et al. (2022), but our study additionally finds that spatial sampling bias alone can be sufficient to produce statistically significant results or distort estimates of methodological biases in many cases when there is no relationship between the environment and habitat suitability. Additionally we present a method for adjusting these tests based on a spatial bias estimate rather than an overall measure of spatial autocorrelation, which can currently be implemented within the ENMTools R package with only slight modifications to normal workflows. We further demonstrate that this method reduces the inflated type I errors associated with previous permutation tests to levels similar to those when using unbiased data. We expect that there are both advantages and disadvantages to this approach when compared to that of Osborne et al. (2022). In particular, the method presented here requires an estimate of spatial sampling bias for the target species over the study region. This is both a strength and a weakness; when the bias estimate is accurate, the pseudoreplicates constructed by sampling from this bias estimate will directly reflect the undesirable effects of spatial sampling bias on the empirical data rather than the overall level of spatial autocorrelation. However, as outlined above, accurately estimating spatial sampling bias is itself inherently difficult and fraught with both practical and conceptual issues that must be dealt with on a study-by-study basis, and the accuracy of the resulting bias estimates is difficult to assess.

Ideally a method for correcting hypothesis tests would include all sources of spatial autocorrelation in the pseudoreplicate simulations that is produced by non-independence of habitat selection by individuals and of data collection by researchers, while leaving intact the spatial autocorrelation that is due to individuals responding independently to a spatially autocorrelated environment. Neither the Osborne et al. (2022) method nor this one are capable of guaranteeing that outcome, and it is possible that no such method may ever be generally available. As such we suggest treating the two approaches as complementary, ideally examining and contrasting the results of each.

More broadly, we feel that these results demonstrate both the value of randomization tests in ENM and the challenges of implementing them correctly. The appeal of big data approaches to ecological questions is that, properly conducted, these studies allow us to understand processes that are operating at a scale that is impossible to manipulate experimentally. Randomization tests are a key component of that, as they allow us to conduct hypothesis tests and measure statistical distributions that we cannot specify a prior, just by repeatedly simulating some aspects of the data. However, proper design of randomization tests is both as important and as complex as the proper design of ecological experiments; careful consideration must be given to what is allowed to vary, what is held constant, and how these changes may affect the resulting inferences.

